# The impact of high-fat, obesogenic diets on brain volume in a commercially available mouse model of fatty liver disease

**DOI:** 10.1101/2024.09.15.613120

**Authors:** Li Jiang, Salaheldeen Elsaid, Cindy Zhan, Xin Li, Su Xu, Sui Seng Tee

## Abstract

The obesity pandemic poses significant health challenges, despite recent advancements in weight loss medications. Mouse models fed obesogenic diets serve as invaluable tools for dissecting the pleiotropic mechanisms underlying weight gain. Here, we utilize these models to analyze brain morphometrics using MRI techniques, inspired by similar findings in human studies linking obesity to brain volume changes. We hypothesize that the mouse model of obesity will exhibit brain volume alterations akin to those observed in obese humans, potentially shedding light on the neurological implications of obesity. To test our hypothesis, mice were provided free access to either regular chow or a diet consisting of high fat and high sugar and MRI scans for total brain volumes as well as volumes of specific brain regions were estimated and compared between obese and control mice. We found that obesogenic diets resulted in ∼13% greater weight gain compared to control chow diets. MRI brain scans revealed reduced total brain volume in obese mice that trended towards significance. In contrast, analysis of specific brain volumes showed an increase in neocortical regions of obese mice, that were significant when compared to controls. In conclusion, diet-induced obesity mouse models are a readily available avatar for studying the obesity epidemic, with significant increases in body weight within a reasonable timeframe. While weight gain among individual mice fed obesogenic diets showed some variability, MRI brain scans were able to reveal significant differences, especially within different anatomical regions of the brain.

## 1. Introduction

The obesity pandemic is well underway, with mounting evidence that weight gain in developed countries continuing to rise^1^. Unfortunately, obesity is a multi-factorial disease, with both genetic and environmental factors reciprocally affecting the outcome of weight gain^2^. While recent high-profile reports of ‘triple receptor’ agonists have resulted in substantial reductions in body weight, a significant proportion of patients do not respond to these medications. The reasons for non-response remain unknown^3^. As such, mouse models of obesity^4^ remain in an invaluable tool to disentangle the intrinsic, genetic mechanisms of unhealthy weight gain, with that of external, environmental factors as we continue to innovate on strategies to combat disorders related to weight gain.

The availability of off-the-shelf models of mice fed obesogenic diets is a valuable resource to define nutrient-based features of obesity^5^. These mouse models are usually derived from inbred, genetically identical mice, fed diets with pre-determined caloric contributions from specific nutrient groups. Therefore, these models serve as avatars to define the effects of particular nutrients on the progression of metabolic disease, independent of genetic background. Furthermore, the wide distribution and availability of commercial disease models^6^ allows a certain degree of consistency, to combat against issues of reproducibility that have plagued the biomedical community^7^.

In this study, we undertake a morphometric analysis of the brains of a commercially available mouse model of obesity. Here, we use widely available magnetic resonance imaging (MRI) techniques to measure a) global brain volume and b) volume of specific brain regions, in mice fed obesogenic diets compared to chow-fed controls. This study design is motivated by reports of volumetric brain volume changes in human subjects, using similar MRI techniques. For example, a recent study of over 12,000 participants from the UK Biobank studies found that obesity is associated with smaller subcortical gray matter volumes^8^. Therefore, we hypothesize that a mouse model of obesity will yield similar changes in brain volume.

## 2. Materials and Methods

### 2.1 Animals

C57BL/6NTac male mice were purchased from Taconic Bioscience (La Jolla, CA, USA). A total of 20 Nonalcoholic steatohepatitis (NASH) and 15 control C57BL/6NTac male mice were included in this study. All mice were maintained on a 12/12-hour light/dark cycle with bedding and cage enrichment, free access to food and water, and in a temperature-controlled environment (22°C). NASH mice were fed a high-fat, high-fructose diet (Research Diet #D09100310, containing 40% kcal from fat, 22% kcal from fructose, and 2% cholesterol) starting at 6 weeks of age and housed at reduced density. Control mice were housed in the same location, also at reduced density, and fed the NIH-31M chow diet (5% kcal from fat).

After 8 weeks on their respective diets (14 weeks of age), all mice were shipped, and group housed (5 per cage) at the University of Maryland School of Medicine in an Association for Assessment and Accreditation of Laboratory Animal Care International-accredited facility. NASH mice continued the research diet, while control mice were fed the regular chow diet (LabDiet #5053, 5% dietary fat, 3.42 kcal/g). All research procedures were conducted in compliance with National Institutes of Health guidelines for animal care and were approved by the Institutional Animal Care and Use Committee at the University of Maryland School of Medicine. The weights of all mice were measured and recorded at the day when the animals arrived and at the day of MRI experiment.

### 2.2 MRI Data Acquisition

At 14 weeks of age, all mice were subjected to MRI brain scans. A total of 5 mice were scanned per day, in an interleaved manner, to prevent batch effects. MR scans were conducted using a horizontal bore Bruker Biospec 7T 70/30 MR Scanner (Bruker Biospin MRI GmbH, Germany) and a Bruker Paravision 6.0 console. Aa Bruker 72mm linear-volume coiland a Bruker ^1^H four-element surface coil array served as transmitter and receiver. Each mouse was intraperitoneal injected with dexmedetomidine (0.03 mg/kg) immediately prior to imaging, and a 0.3% isoflurane concentration was maintained. Animal respiration rate and body temperature were monitored using a MR-compatible small-animal monitoring and gating system (SA Instruments, Inc., New York, USA). The animals’ body temperature was maintained at 37–38.5°C by circulating warm water in a bath.

Structural T2-weighted (T2-weighted) images covering the whole brain were obtained using a 2D Rapid Acquisition with Relaxation Enhancement (RARE) pulse sequence along the coronal direction. The imaging parameters are the following: Repetition Time (TR) = 2500ms, echo time (TE) = 30ms, number of averages = 6, flip angle = 90°, Field of view 150 voxels, in-plane resolution was 0.12mm x 0.12 mm, slice thickness = 0.5 mm with no gap between slices, and total number of slices = 26.

### 2.3 MRI Data Processing and Analysis

#### 2.3.1 Study-specific T2-weighted brain template

**Figure 1:**
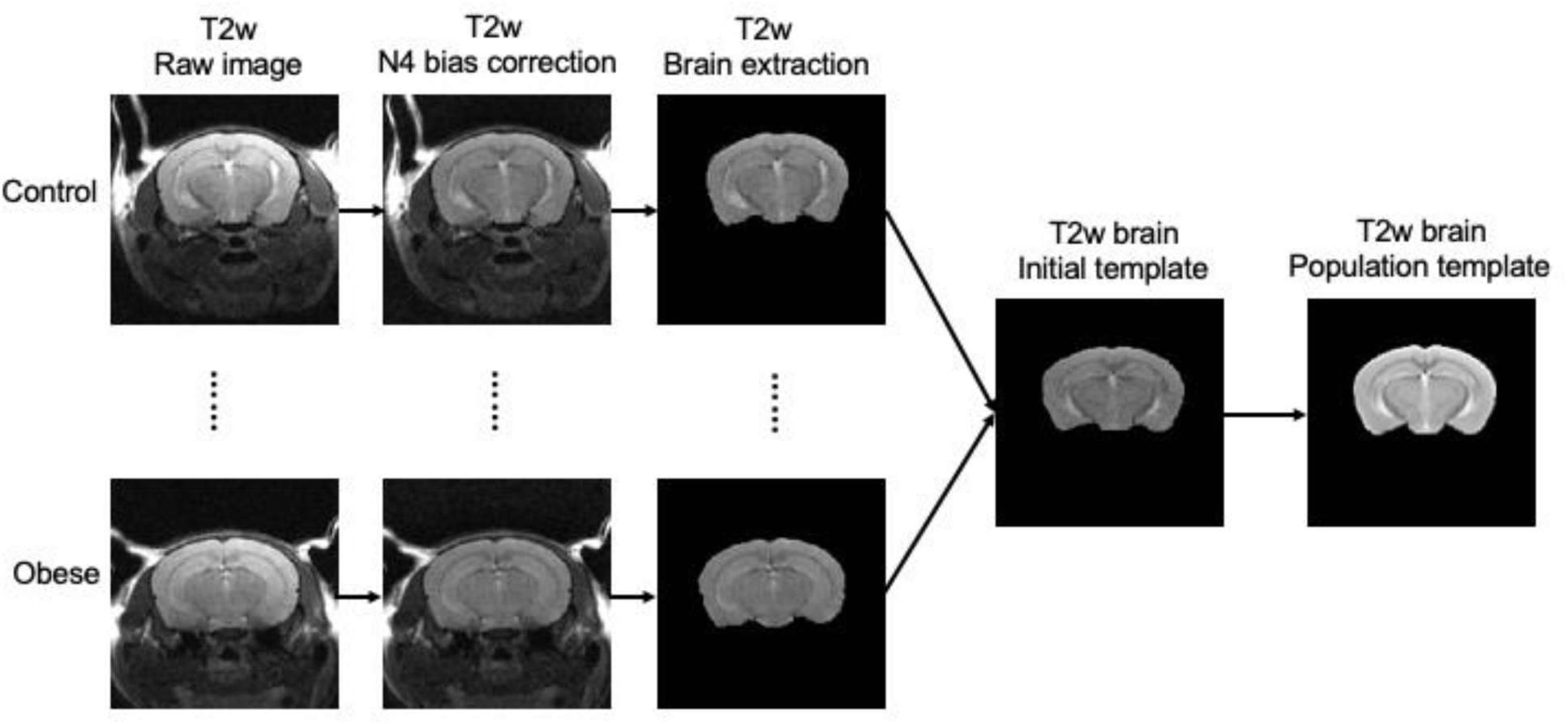
Schematic of image processing procedures to derive whole brain templates for volumetric analysis. Briefly, raw T2 weighted (T2-weighted) images were subjected to intensity inhomogeneity correction (N4 bias correction), before segmentation to extract brain-only regions. These preprocessed, skull-stripped images (n = 15 from control and n = 15 from obese) were randomly selected to create a study-specific T2-weighted brain template.

To register all individual T2-weighted image into a common space, a study-specific T2-weighted template image was generated and the steps were demonstrated in Figure 1.

First, all T2-weighted images from the Control group were collected for template generation, and the image with the best orientation and quality was chosen as the initial template. The selected T2-weighted images were then bias-corrected using the “N4BiasFieldCorrection” tool from ANTs package^9^ and brain extracted using the 3dAutomask from AFNI software package. The resulting brain masks were visually checked and manually modified using ITK-SNAP tool. Subsequently, the “buildtemplateparallel.sh” script from ANTs was used to create the T2-weighted template based on the T2-weighted brain images and the initial template. The involved steps involved an iterative approach, mutual information as a similarity metric for registration, and symmetric normalization transformation through diffeomorphic warping techniques.

#### 2.3.2 T2-weighted image processing and brain volume estimation

As above, all T2-weighted images from Control and NASH mice groups were bias-corrected and brain masks were extracted and manually modified. The resulted T2-weighted brain images were then registered to the T2-weighted brain template using affine transformation. We performed three different strategies to investigate the whole brain volume and regional volumetric difference between the Control and NASH groups.

##### Whole brain volume Analysis

Upon visually inspecting the registered T2-weighted brain images, we observed that not all T2-weighted images covered the entire brain. Therefore, we restricted our analysis to coronal slices 3 through 22, moving from caudal to rostral regions, to ensure that brain volume measurements were consistent across all animals. AFNI’s 3dAutomask was applied to the registered T2-weighted brain images to generate brain masks. These masks were visually inspected, and manual edits were made when necessary to ensure accuracy. The brain volume was estimated by the total number of voxels within the extracted brain mask multiplied by the voxel size.

##### Segmented Coronal Volume Analysis

We employed a Segmented Coronal Volume Analysis method, where coronal slices were divided into three regions: posterior (slices 3-7), middle (slices 8-19), and frontal (slices 20-22) as illustrated in Fig 3A. The volumes within these segmented regions were calculated to provide a more detailed assessment of brain volume distribution.

##### ROI volume Analysis

We conducted ROI-based volume analysis to investigate the specific volume differences between control and NASH groups. First, six ROIs were manually delineated on our study-specific T2-weighted brain template, using the Allen Mouse Atlas as a reference. These ROIs included key brain regions: the cerebellum, cerebral cortex/neocortex, hippocampus, thalamus, caudate-putamen (CPU), and prefrontal cortex, as illustrated in Fig. 4A. To obtain the corresponding ROIs for each individual animal, we performed deformable registration of the individual T2-weighted brain images to the study-specific template. The inverse transformation matrix was then applied to the template ROIs to project them back into the individual brain space. To ensure accuracy, the ROI masks in individual space were visually inspected, and manual corrections were made as necessary. Finally, the volumes of the ROIs were estimated. We implemented an adjustment procedure to account for variations in whole brain volume by normalizing the ROI volumes. This was done by calculating the ratio of each ROI’s volume to the corresponding whole brain volume.

### 2.4 Statistical Analysis

To assess differences in the volumetric measurements between the control and NASH mouse groups, we conducted a two-sided two-sample t-test. This statistical analysis was performed to identify statistically significant differences between the two groups.

### 2.5 Correlation between Brain Volumes and Body Weights

We employed Pearson correlation analysis to explore the relationships between brain volumetric measurements and body weight. The goal was to identify potential associations between these variables. Significant correlations were determined based on a predefined significance level of p-value less than 0.05.

## 3. Results

### 3.1 Obesogenic Diet Results in Significant Increase in Body Weight of Mice

After 8 weeks on an obesogenic diet high in fat and fructose, mice showed noticeable differences in appearance compared to chow-fed controls. The most prominent change was oily or greasy fur, consistent with observations in other high-fat diet models. Truncal obesity was also evident, as shown in Fig. 2. As expected, the body weights of obese mice were significantly higher than those of the chow-fed controls. At 14 weeks of age (corresponding to 8 weeks of dietary intervention), obese mice weighed approximately 13% more than chow-fed mice. Despite this, there was greater variability in body weights among the obese mice, with a mean weight of 32.7 g and a modal value of 36 g. In contrast, control mice had an average weight of 28.9 g, with most weighing between 28 g and 29 g. Notably, all mice are of a similar genetic background and are group-housed (5 per cage), so the source of the weight variability in obese mice remains unclear.

**Figure 2:**
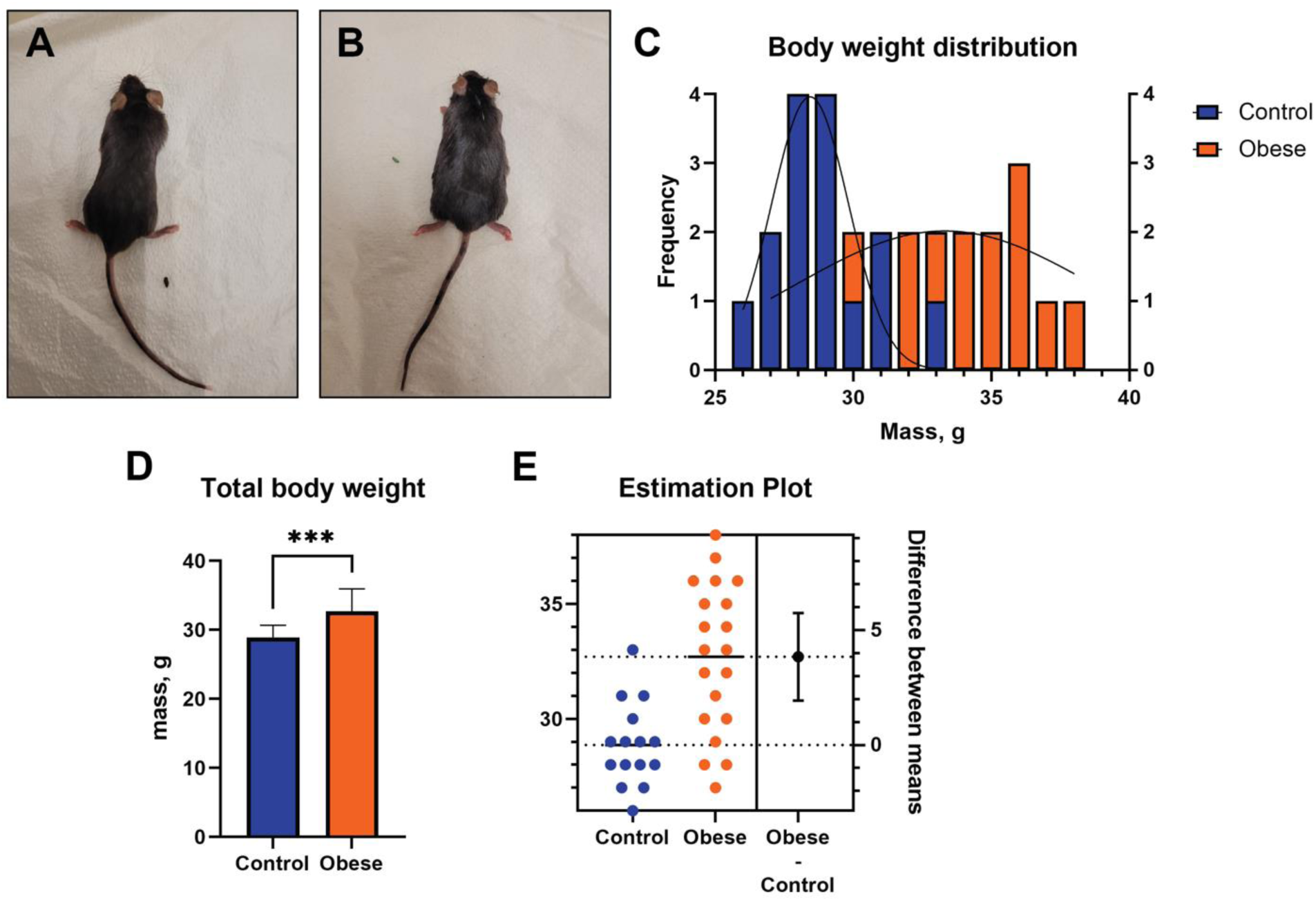
Representative photographs of male (A) chow-fed control or (B) obesogenic, high fat, high fructose-fed mice. (C) Distribution of body weights in control and obese mice (control n = 15, obese n = 25). (D) Average total body weights for control mice were 28.9 ± 1.8g vs. obese mice at 32.7 ± 3.3g (all values are mean ± SEM). (E) Estimation plot of the effect size of dietary intervention on control vs. obese mice. The difference between means of obese and control populations is 3.8 ± 0.9g, with a 95% confidence interval between 1.9 and 5.7g. Thus, the difference between control vs. obese body weight is statistically significant (p < 0.05, two-tailed two-sided two-sample t-test).

### 3.2 Brain Volumetric Results Compared between Obese Mice vs. Controls

#### Whole brain volume

Figure 3A illustrates the brain slices used for estimating whole brain volume. As shown in Fig. 3B, the average whole brain volume of Control group is 645.02 ± 12.27 mm^3^ and NASH mice group is 642.21 ± 9.62 mm^3^. The NASH mice had reduced whole brain volume but not significant (T = −0.760, p = 0.4524).

**Figure 3:**
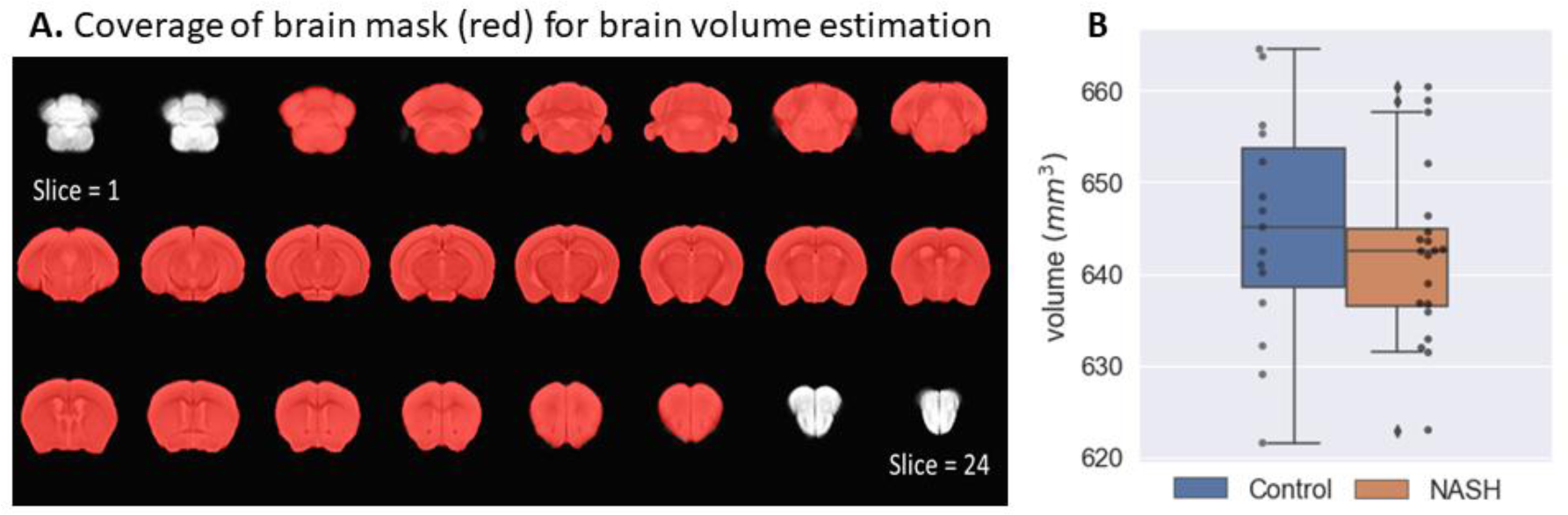
(A) Representative images of a coronal sections with slices selected for whole brain volume analysis highlighted in red. (B) Two-sample two-sided t-test between Control and NASH groups.

#### Segmental brain volumes

Figure 4A illustrates the subdivision of the brain into three segments. As shown in Fig. 4B. NASH mice showed significant decrease in posterior area (T=-2.947, p = 0.0058) and frontal area volumes (T = −2.374, p = 0.023) and an increase but not significant in the middle area volume (T = 0.624, p = 0.5365) in the NASH group.

**Figure 4:**
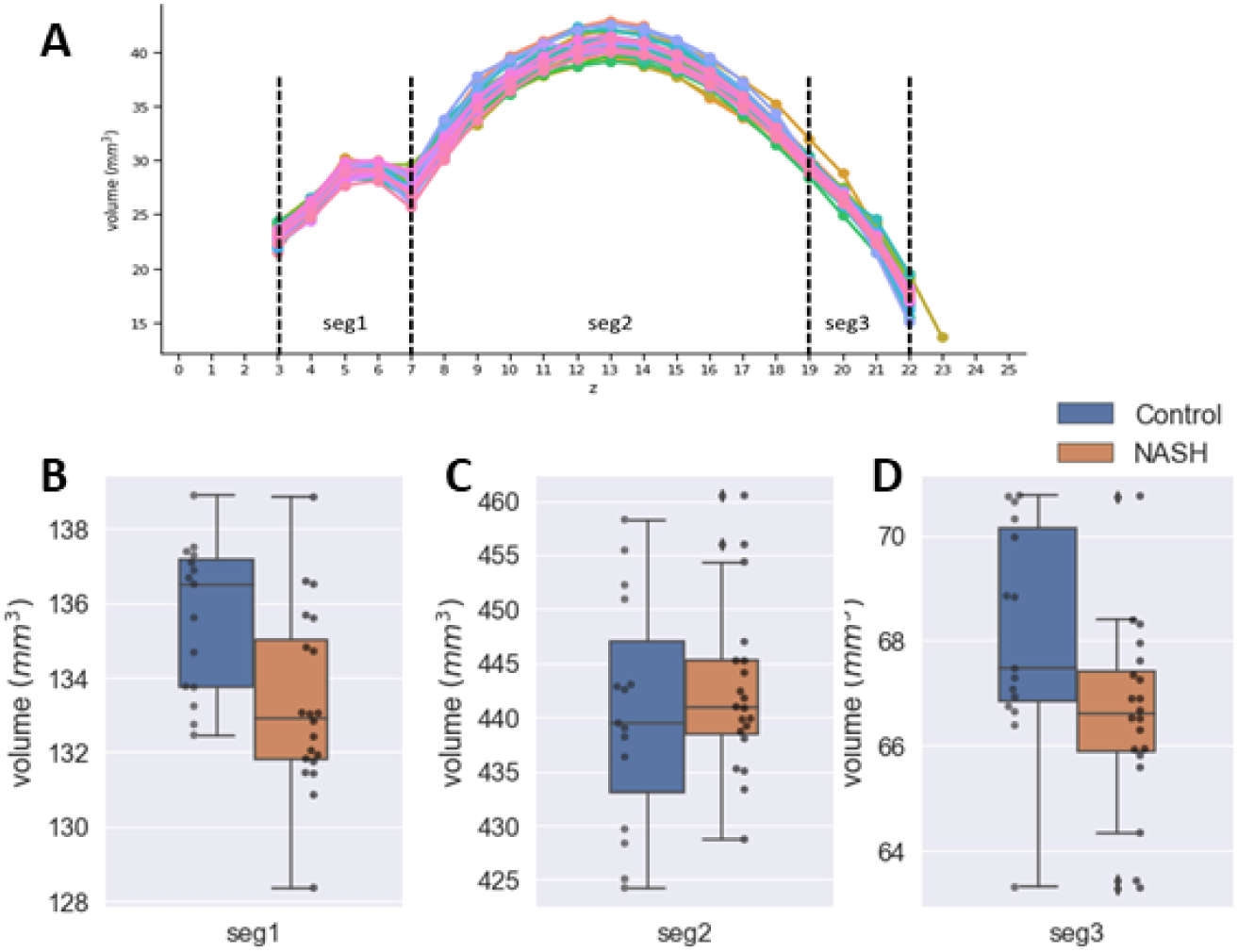
(A) Segmental volumetric analysis, with brain regions divided in three sections: posterior (slices 3-7, labeled Seg1), mid-brain (slices 8-19, labeled Seg2) and frontal (slices 20-22, labeled Seg3). (B-D) Comparison of segmental volumes between Control and NASH groups. Black dots in (B-D) represent the individual volumes for each segment. Volumes of Seg 1 and Seg3 in obese mice are reduced in obese mice (p = 0.0058 and p = 0.023, respectively). Seg2 brain volumes trended higher in obese mice vs. controls, but this change was not statistically significant (p = 0.624).

#### ROI brain regional volumes

Fig. 5A illustrates the brain subregions for which we estimated the volumes. As shown in Fig. 5B, we found that the volume of the neocortex in obese animals were significantly larger than that of controls. In contrast, no other region displayed a statistically significant difference in terms of regional volumes between the two groups.

**Figure 5:**
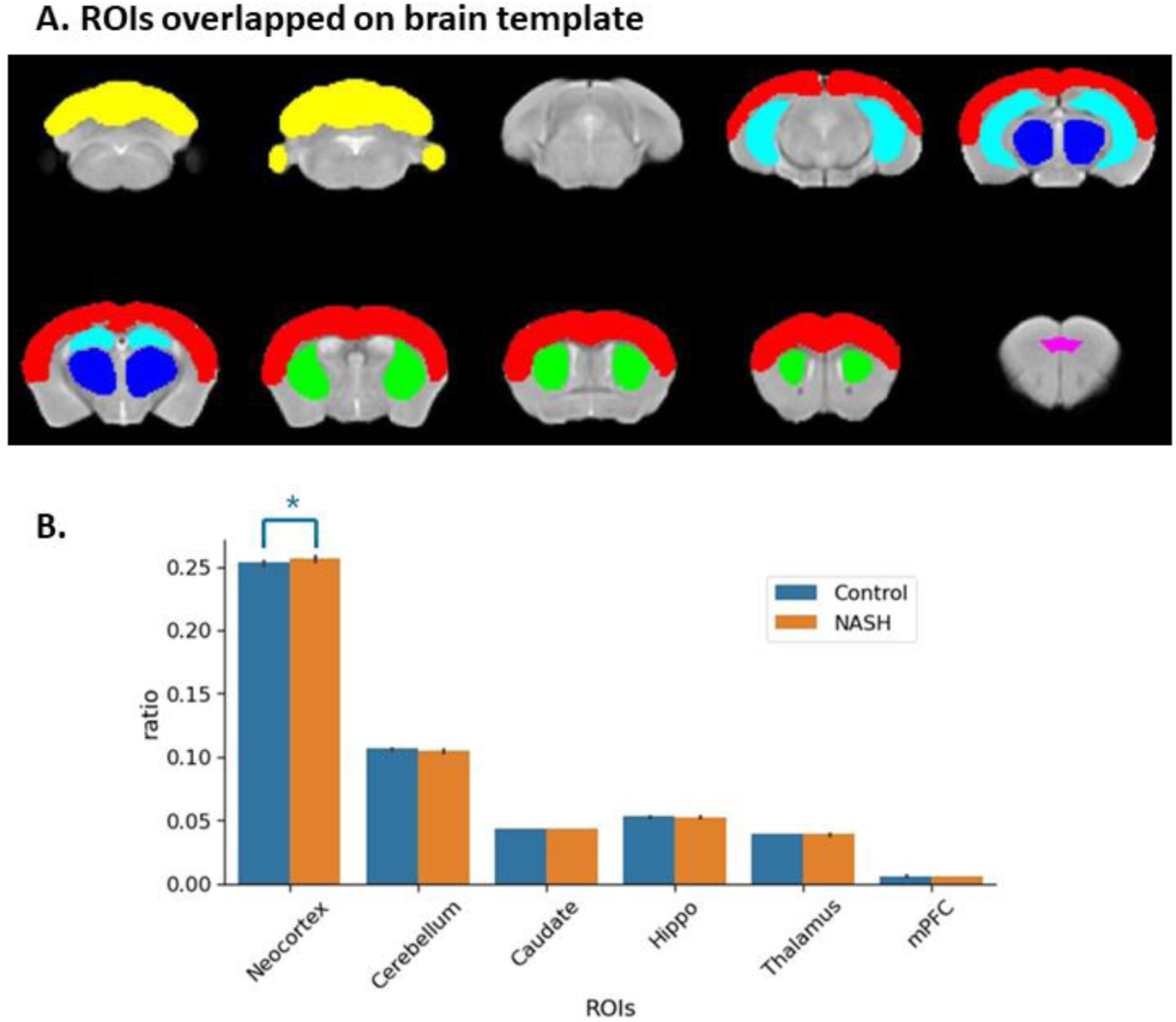
(A) Representative regions of interest (ROI). These include cerebellum (yellow), neocortex (red), caudate putamen (CPU, green), thalamus (blue), hippocampus (light blue) and middle prefrontal cortex (mPFC) (magenda). (B) Ratio metric volume quantification of different brain regions, where a specific regional volume was expressed as a ratio of total brain volume. The neocortical volumes of obese mice were significantly larger than control mice. In contrast, there was no statistically significant volumetric changes in the cerebellum, CPU, hippocampus, thalamus and mPFC.

### 3.3 Correlation between Brain Volumetric Measures and Body Weights

We performed Pearson correlation between brain volumetric measures and body weights using SPSS version 26. The results showed that significant correlation only occurs between neocortex volume and body weight (Figure 6 and supplemental Table2). The control group showed significantly negative correlation between neocortex volume and body weight (r = −5.49, p = 0.034), while NASH group only showed a negative trend but not significant (r=-0.015, p=0.949). When we combined the Control and NASH mice together, a strong positive correlation was observed (r = 0.486, p = 0.003). No significant correlation was found in whole brain and other regional brain volumes.

**Figure 6.**
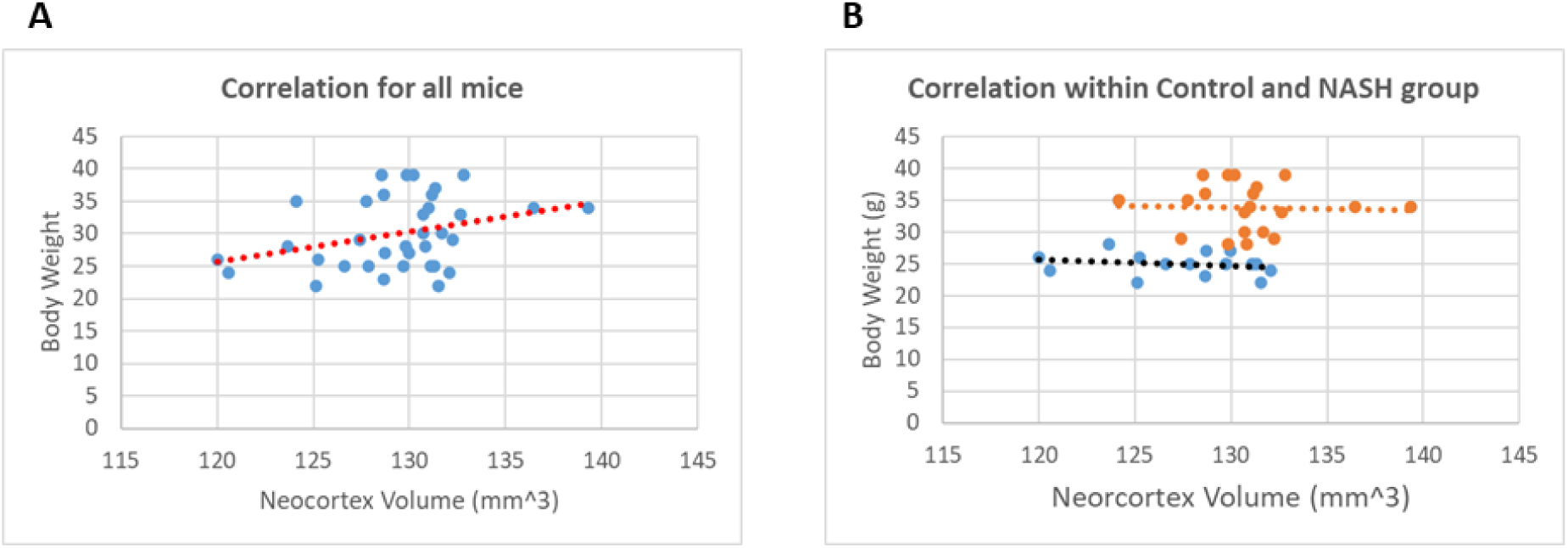
Pearson Correlation between neocortex volume and body weight. (A) for all mice; (B) control group and NASH group respectively. The Control group shows a strong negative correlation, while the NASH group does not show as pronounced a trend. The combined data might reflect the overall range of body weights and brain volumes, creating a positive correlation due to the larger spread of values.

## 4. Discussion

In this study, we examined the relationship between brain volumes and obesity, in mice fed a high-fat, high fructose diet. The motivation of using inbred, male mice allows investigation of the effect of overnutrition on the brain, independent of genetic and gender-based confounding factors. In this model of diet-induced obesity, we measured differences in total brain volumes, as well as volumes in specific regions of the mouse brain.

Based on large-scale studies in humans, we tested the hypothesis that obesity in mice is associated with a decrease in cortical volumes^8^. However, we did not find support for this hypothesis in our experiments. In fact, we report an increase in neocortical volumes in obese mice. In contrast, other regions of the brain showed no significant difference in volume. Similarly, total brain volumes were not significantly different between control and obese mice. It was only when brain volumes were divided into 3 different sections (posterior vs. mid-brain vs. frontal), that a difference was seen where posterior and frontal brain volumes in obese mice were significantly lower than controls.

The inconsistencies between our rodent study vs. human studies is not without precedent. Indeed, a recent publication noted the divergence of MRI-based volumetric scans in patient-based studies. In this report, the authors focused on grey matter volumes and body weight, summarizing 21 studies performed between 2008 and 2016^12^. The authors noted the inconsistencies of volumetric changes reported in overweight populations, where the majority of studies reported decreases in grey matter, but the location of volumetric change varied widely. Furthermore, several studies reported increases in regional grey matter volume. Several reasons were posited as driving factors behind these inconsistencies. Chiefly, obesity is driven by complex interactions between the environment^13^, behavior^14^ and individual genetic profiles^15^. Additionally, many patient studies in the clinic utilize body mass index (BMI) as a surrogate for obesity^16^. However, while BMI represents a simple and low-cost measure for assessing obesity, the index itself represents an amalgamation of multiple metabolic disorders, lacking the sensitivity to detect specific causes of obesity^17^. These shortcomings directly motivate our approach in using mouse models of diet-based obesity, to provide a diet-centric, volumetric signature of the obese brain, to evaluate the contribution of unhealthy diets to structural brain changes.

While there is a wealth of volumetric analysis of the human brain in metabolic disorders, there are fewer examples of corresponding studies in animal models^18^. For example, female mice exposed to diets containing different levels of fat failed to show differences in total brain volume^19^, consistent with results presented in our study. It is worth noting that the referenced study utilized T2-weighted MRI of *ex vivo*, fixed mouse brains after euthanasia, as opposed to our study in live mice. A more recent study attempted volumetric analysis in live mice exposed to high-fat diets^20^. Here, the diet-induced animal model of non-alcoholic fatty liver disease (DIAMOND) mouse model was used. This model is an isogenic cross between the C57BL/6J mouse strain, with the 129S1/SvImJ strain, obtained through repeated brother-sister mating over a four-year time frame^21^. Here, the authors report a statistically significant reduction in total cerebral brain volume (TCBV), starting from 4 weeks post-dietary intervention, that persists for the next 24 weeks of obesogenic diet feeding^20^. The reduction in TCBV reported is in stark contrast to our study, where we noted an increase in neocortical volumes in obese mice. Several reasons are offered for this inconsistency, highlighting the relative strengths and weaknesses of our study.

Firstly, while the majority of studies in mice utilize inbred rodents with ‘stable’ genetic backgrounds, there is still a difference in mouse strains. In our study, the C57BL/6NTac sub-strain originates from the original C57BL/6 strain established in the 1920s. This parent strain subsequently diverged to C57BL/6J and C57BL/6N, from the Jackson Laboratories and the National Institutes of Health (NIH) respectively^22^. The C57BL/6NTac was generated in 1991^23^. Here, we argue that the wide availability and defined origins of the sub-strain used in our studies promote future attempts to replicate this work, to assess potential areas of inconsistency. In contrast, the DIAMOND mouse model used to be a commercial product, but to the best of our knowledge, is now currently unavailable for purchase.

Secondly, the formulation of obesogenic diets used in mouse models of metabolic dysfunction represents a source of potential inconsistency. The mice used in this study were fed a well-defined, purified diet with 40 kcal% fat, 22% fructose and 2% cholesterol^24^. This diet is widely used for its ability to induce stepwise progression of non-alcoholic fatty liver disease (NAFLD), from simple steatosis to steatohepatitis with fibrosis and finally cirrhosis. Beyond liver dysfunction, mice demonstrated increased total cholesterol, heightened fasting insulin and lower adiponectin, mirroring the multi-faceted presentation of metabolic syndrome in humans^25^. As a comparison, the DIAMOND mouse model of obesity used a diet containing 42 kcal% fat, 0.1% sucrose and a sugar water solution containing 23.1 g/L fructose and 18.9 g/L glucose. While both diets result in a significant increase in body weight, alterations of specific nutrients result in appreciable phenotypic differences. As an example, the use of non-*trans* containing fat worsened insulin resistance, when all other dietary and strain-specific differences were kept constant^26^. Thus, it is possible that differences in brain volumes seen in our study vis-à-vis other published research stems from the nutritional composition of the diets used. However, it is worth noting that that dietary formulation that was used in this study is also commercially available, allowing for attempts at reproducing this study in the future if desired.

Finally, the methodology used in our study to measure brain volumes may be a cause for inconsistency. Here, we utilized a T2-weighted imaging sequence to acquire images. In contrast, the DIAMOND mouse study utilized a T1w imaging sequence^21^. T1w sequences represent the recovery time of the longitudinal magnetization, after excitation with a radiofrequency (RF) pulse. In contrast, T2-weighted sequences emphasize the recovery of transverse magnetization after application of an RF pulse^27^. T1 and T2 relaxation times are dependent on the average contributions from the chemical environment that a water proton experiences. Typically, T1w images have lower contrast than T2-weighted equivalents, resulting in many T1w imaging studies requiring administration of a contrast agents such as gadolinium^28^. This motivated our choice of utilizing T2-weighted imaging for volumetric determination, due to its excellent reputation for providing good soft-tissue contrast^29^. Therefore, differences in MRI methodology may also result in the inconsistency seen between this volumetric study compared to earlier published work.

In summary, this study demonstrated the ability to use commercially available mouse models of obesity for studying the relationship between diet and brain volume. When male C57BL/6NTac mice are challenged with a diet high in fat, fructose and cholesterol, for a total of 8 weeks, we report a statistically significant increase neocortical volume. While this volumetric difference remains to be validated in other models of overnutrition, the study design presented here facilitates robust reproducibility studies based on the findings reported here. Future studies will focus on the reversibility of this volumetric phenotype, especially with interventions that reduce body weight, such as physical activity or hormone receptor agonists.

## Declarations

Ethics Statement: All animal experiments were performed after approval by the University of Maryland Baltimore IACUC committee (Protocol #0822007) prior to conducting the research. The authors confirm that these experiments were conducted according to established animal welfare guidelines the American Veterinary Medical Association (AVMA) Guidelines.

