## Supplemental Tables for "The impact of high-fat, obesogenic diets on brain volume in a commercially available mouse model of fatty liver disease"

**Table 1. Two-sample T-test of ROI-based regional brain volumes between Control and NASH groups**.

|  | **Controls** | | **NASH** | | **t-test** | |
| --- | --- | --- | --- | --- | --- | --- |
| **ROI** | **mean** | **std** | **mean** | **std** | **T value** | **p value** |
| **Neocortex** | 0.199 | 0.004 | 0.205 | 0.003 | 4.993 | < 0.001 |
| **Cerebellum** | 0.107 | 0.002 | 0.107 | 0.002 | 0.093 | 0.926 |
| **Caudate** | 0.044 | 0.001 | 0.044 | 0.001 | 1.480 | 0.148 |
| **Hippocampus** | 0.053 | 0.001 | 0.054 | 0.001 | 1.290 | 0.206 |
| **Thalamus** | 0.040 | 0.001 | 0.040 | 0.001 | 0.720 | 0.477 |
| **Middle Prefrontal Cortex** | 0.006 | 0.000 | 0.006 | 0.000 | -0.780 | 0.441 |

**Table 2. Pearson correlation between ROI-based regional brain volumes and body weights.**

|  |  | **All Mice** | **Control** | **NASH** |
| --- | --- | --- | --- | --- |
| **wholebrain** | Pearson Correlation | -0.056 | 0.211 | -0.052 |
|  | Sig. (2-tailed) | 0.749 | 0.451 | 0.826 |
|  | N | 35 | 15 | 20 |
| **Neocortex** | Pearson Correlation | .486** | -.549^*^ | -0.015 |
|  | Sig. (2-tailed) | 0.003 | 0.034 | 0.949 |
|  | N | 35 | 15 | 20 |
| **cerebellum** | Pearson Correlation | -0.154 | -0.034 | -0.297 |
|  | Sig. (2-tailed) | 0.378 | 0.905 | 0.204 |
|  | N | 35 | 15 | 20 |
| **Caudate** | Pearson Correlation | 0.133 | -0.412 | -0.038 |
|  | Sig. (2-tailed) | 0.446 | 0.127 | 0.875 |
|  | N | 35 | 15 | 20 |
| **Thalamus** | Pearson Correlation | 0.039 | -0.395 | 0.000 |
|  | Sig. (2-tailed) | 0.825 | 0.146 | 1.000 |
|  | N | 35 | 15 | 20 |
| **Hippocampus** | Pearson Correlation | 0.072 | -0.308 | -0.142 |
|  | Sig. (2-tailed) | 0.681 | 0.263 | 0.550 |
|  | N | 35 | 15 | 20 |
| **mPFC** | Pearson Correlation | -0.123 | -0.010 | -0.001 |
|  | Sig. (2-tailed) | 0.480 | 0.973 | 0.995 |
|  | N | 35 | 15 | 20 |
